# Transgenerational coexistence history attenuates negative direct interactions and strengthens facilitation

**DOI:** 10.1101/2023.02.08.527660

**Authors:** Anja Schmutz, Christian Schöb

## Abstract

**Background:** Interactions among species are a fundamental aspect of biodiversity and driving ecosystem functioning and services. Species interactions include direct (pairwise) interactions among two species and indirect interactions that occur when a third species interacts with the two others and changes the direct interactions between the two. In a three-species interaction network, these interactions can be transitive (where one species outperforms all others) or intransitive (where each species outperforms another). How direct and indirect interactions influence ecosystem functions in crop systems, and how diversification and evolutionary adaptation can influence those interactions and therefore ecosystem functions has not been studied.

**Methods:** A common garden experiment was conducted with crop communities in monocultures, 2- and 3-species mixtures that had either a common or no coexistence history (i.e. community adaptation) for three years. Net, direct and indirect interaction intensities were estimated and compared between the diversity levels and coexistence histories. Furthermore, species interaction networks were inspected for transitive/intransitive interactions.

**Results:** We found evidence for lower competition in mixtures and for reduced negative direct interaction intensity and enhance facilitative effects upon community adaptation. We could further show that indirect interactions were generally less important for community adaptation than direct interactions. Additionally, we showed that community adaptation has the potential to shift interactions in the species interaction networks from competitive intransitive into pairwise competitive interactions where interactions occurred mainly between two species.

**Synthesis:** Co-adapted crop species with reduced negative interactions might have the potential to enhance productivity especially in more diverse cropping systems. This supports the notion that intercropping is a vital part towards a more sustainable agriculture and one with further yield potential when developing cultivars adapted to grow in mixtures.

## Introduction

Interactions among species are omnipresent. Many of these interactions are competitive, where species compete for certain resources (Levine, 1976). Nevertheless, interactions among species can also be positive (i.e. facilitation) and occur when one species enhances the fitness of another species (Brooker et al., 2008; Hunter and Aarssen, 1988). These (positive and negative) interactions are important for community assembly (Schöb et al., 2013), species coexistence (Levine et al., 2017) and can have evolutionary consequences for the species involved (Thorpe et al., 2011).

In a community with two individuals, the direct interactions between these two individuals are often very intense (**Fig. 1a**). This direct interactions are usually attenuated in communities with more than two individuals (Aschehoug and Callaway, 2015; Levine et al., 2017) due to the presence of indirect interactions which attenuate the competitive effects of the individuals involved in a pairwise interaction (**Fig. 1b**) (Callaway, 1997; Miller, 1994). Specific forms of interaction networks involving direct and indirect interactions are intransitive networks (Allesina and Levine, 2011; Gallien et al., 2017), where each species is more competitive than another species in the community (**Fig. 1c**), thereby promoting coexistence. Alternatively, transitive networks show a hierarchical structure in competitive ability (**Fig. 1d**), which tend to result in the most competitive species becoming dominant and the less competitive species being competitively excluded (Hardin, 1960). Hence, direct and indirect interactions and their interaction network have ecological and evolutionary consequences on species communities (Lawlor, 1979; Schöb et al., 2013; Strauss, 1991; Wootton, 1994).

**Fig. 1.**
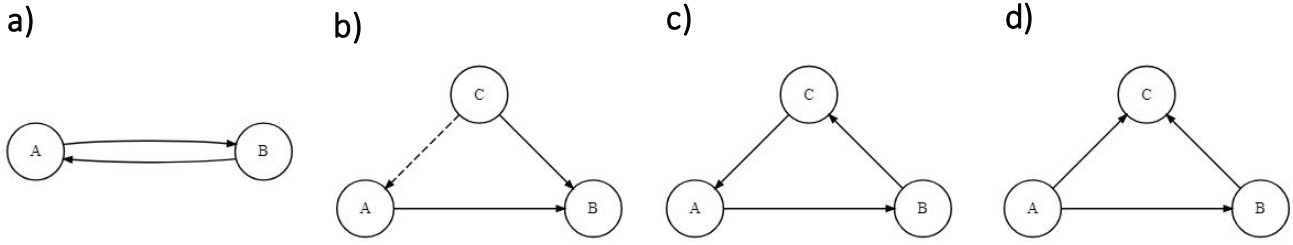
Different type of interactions between two or three individuals. (**a**) Direct (pairwise) interaction with reciprocal negative effects between A and B. (**b**) Indirect interaction when C interacts with A and B. The original negative interaction from B on A is attenuated (or disintegrated) as C is more competitive against B. Hence, C has a positive indirect effect on A (dashed line). (**c**) Intransitive interaction between A, B and C where each individual is more competitive than another one (“rock-paper-scissor” game). (**d**) Transitive (“hierarchical”) interaction between A, B and C where A is more competitive than B and C and B more competitive than C.

In an crop system, the reduction of negative interactions between individuals is essential to mitigate negative effects on fitness or yield of the individuals composing the crop community (Donald, 1981; Vandermeer, 1989; Weiner et al., 2017). However, the commercial crop species used nowadays are highly bred which might have selected for high-yielding “selfish” individuals that intensify competition (Weiner, 2019) rather than promoting cooperative behaviour (Wuest et al., 2022). Hence, studying interactions among species helps to understand how species are competing with others in the community. Furthermore, breeding of crop species might have reduced phenotypic plasticity (Brooker et al., 2022; Vilela and González-Paleo, 2015), although phenotypic plasticity can help plants to reduce competition (Callaway et al., 2003; Schmutz and Schöb, 2022).

Therefore, breeding crops with reduced competition (or enhanced facilitation) and a more cooperative behaviour is a target (Wuest et al., 2022). Indeed, studies in natural plant communities have demonstrated that natural selection in diverse communities can reduce competition, increase facilitation and in consequence increase ecosystem functioning (Schöb et al., 2018; van Moorsel et al., 2018). Similarly, in an annual crop system, crop communities with a common coexistence history showed decreased competition compared to communities composed of crops that did not share a common history (Stefan et al., 2022). This suggests that evolutionary plant breeding of crops in diverse communities (i.e. mixtures) might select for reduced competition, a more cooperative behaviour and more community level yield.

While growing crops in species mixtures tends to show reduced competition and higher yields than monoculture due to the simple fact of niche differences between species (Zuppinger-Dingley et al., 2014), breeding for mixture ecotypes could unfold an additional yield potential of crop mixtures (Bourke et al., 2021; Moore et al., 2022). Using evolutionary processes in mixtures as a tool for breeding mixture ecotypes seems like an easy and straightforward way towards that aim.

In this study, we aimed to link evolutionary adaptation through a transgenerational coexistence history (i.e. community adaptation) with direct and indirect interaction intensities within (i.e. intraspecific) and between crop species (i.e. interspecific). We further examined how community adaptation affects the crop species interaction network. Particularly, the following research questions were addressed: Does community adaptation change intra- and interspecific net, direct and indirect interaction intensities (**question 1**)? Are both direct and indirect interactions equally important in community adaptation (**question 2**)? How does community adaptation change the interaction network towards transitivity or intransitivity (**question 3**)? We hypothesised that community adaptation decreases negative net and direct interactions (i.e. competition), and increases positive indirect interactions (i.e. facilitation) (**question 1**) and that indirect interactions play an important role in community adaptation (**question 2**). Lastly, we expected that upon community adaptation interactions in the species interaction network are significantly shifted and that upon community adaptation interactions are predominantly intransitive (**question 3**). To test these hypotheses, a common garden experiment was conducted with six different crop species that had either a common coexistence history or no coexistence history. The comparison between communities with or without coexistence history allows to investigate how transgenerational coexistence shape interactions among plant individuals. Plants were grown in monocultures (i.e. intraspecific interactions) and mixtures (i.e. interspecific interactions) and net, direct and indirect interactions within and between species were calculated and tested for the hypotheses. Furthermore, species interaction networks with direct and indirect interactions were designed, and the interplay of these interactions were compared.

## Material & Methods

### Experimental design

Plants were grown in an outdoor experimental garden in Torrejón el Rubio, Cáceres, Spain (39°48’48”N 6°00’01”W) from February 2021 until July 2021 in square plots measuring 25×25 cm. These plots were arranged in beds of 4×40 plots, where the cultures were randomly allocated – except the single plants, which were grown in two separate beds to prevent any aboveground interactions. The beds were irrigated through the whole growing season with an automated irrigation system with thresholds that maintained soil moisture between 50% and 75% of field capacity. To minimise interaction with weeds, plots were regularly weeded. Fertiliser was not applied to the plots. For more detail about the experimental design and the experimental garden (e.g. information about the weather) see Schmutz & Schöb (2023).

Six different crop species from three functional groups were grown: the cereals oat (*Avena sativa* var. Previsión) and summer wheat (*Triticum aestivum* var. Cabezorro), the legumes lentil (*Lens culinaris* var. de la Armuña) and lupin (*Lupinus angustifolius* wild type) and the herbs camelina (*Camelina sativa* n.a.) and coriander (*Coriandrum sativum* wild type). These species had either a common coexistence history (seeds from plants grown in mixtures with a specific species composition, *community* selection history) or no coexistence history (seeds from plants grown as single plant, *single* selection history) for three years previous to this experiment. Seeds from *community* selection history were chosen from mixtures as both intra- and interspecific interactions are present. These species and selection histories were grown as single plant (one individual), in 2×2 (four individuals) and in 3×3 Latin squares (nine individuals) (**Table S1**). The two positions within the 2×2 and the three positions with the 3×3 Latin squares were either occupied by the same species (monocultures) or with other species (mixtures) (**Fig. S1**). For the mixtures, all possible combination between species from the different functional groups were planted together. Within one plot, the plants were either from *community* or from *single* selection history (**Table S1**). Plants were harvested after complete senescence and seed maturity (staring 7 June 2021). Aboveground biomass was collected for each individual in each plot and all biomass except seeds was subsequently dried at 80°C for 72h and weighed.

### Data analysis

Data analysis was carried out in R version 4.2.0 (R Core Team, 2022). Mean biomass was calculated from the three individuals that occupied the same position within the 3×3 Latin square (**Fig. S1**). Biomass from single plants and 2×2 Latin squares was only used to calculate interaction intensity in the 3×3 Latin squares. Hence, mean biomass in the 2×2 Latin squares was calculated for each species in each species composition and selection history. Along the same vein, mean biomass in the single plants was calculated for each species and selection history.

To calculate interaction intensities, the relative interaction index (RII) was used (Armas et al., 2004). Net interaction intensity (RII_net_) was calculated for each species in the 3×3 Latin squares.

The corresponding biomass when grown as single plant was used as control (**Eq. 1**). RIInet gives information about the effect of the whole community on a species. Direct (RII_direct_) and indirect interaction intensity (RII_indirect_) were calculated between each position (a, b and c) in each 3×3 Latin square plot separately. Direct interaction was calculated from the biomass of the species that received the interaction (“receiver” species) in the 3×3 Latin square and the biomass of the receiver species in the 2×2 Latin square that was lacking the species that imposes the direct interaction (“donor” species) on the receiver species (**Eq. 2**) (Aschehoug and Callaway, 2015). The indirect interaction was calculated from the product of direct interaction intensity in the 3×3 Latin squares which were mediated by the third species (**Eq. 3**, **Fig. 2**). To prevent pseudoreplication in the monocultures, the mean of direct and indirect interaction for each monoculture plot was calculated. The total interaction intensity (RII_sum_) of the donor species on the receiver species was calculated by the sum of direct and indirect interaction intensities (**Eq. 4**). Direct, indirect and total interaction intensities are indices of interaction strength of the donor species on the receiver species (i.e. interaction strengths between species pairs).

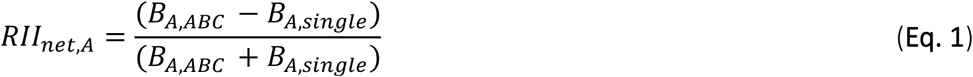

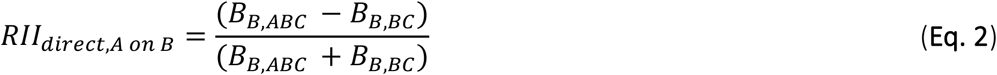

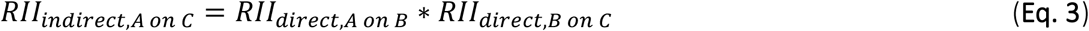

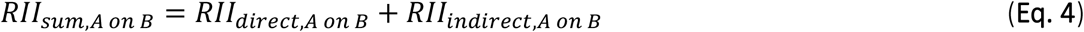

**Fig. 2.**
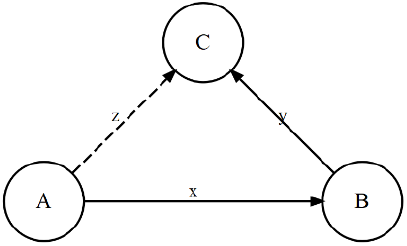
Example of the calculation of direct and indirect interactions. The direct interaction of A on B in Eq. 1 equals x, the indirect interaction of A on C in Eq. equals the product of x and y. In this example, the species B would the receiver of direct interaction from the donor species A (x), the species C would be the receiver of direct interaction from donor species B (y) and species C would be the receiver of indirect interaction from donor species A (z).

Subsequently, linear mixed models (LMM) were conducted for net, direct, indirect and total interaction intensities (R packages *lme4* (Bates et al., 2015) and *lmerTest* (Kuznetsova et al., 2017). For the net interaction intensity, RII_net_ (square-root transformed) was the response variable, species, culture (monoculture vs mixture), selection history (community vs single) and all possible interactions were the explanatory variables, the species composition was the random term. For direct, indirect and total interaction intensities, RII_direct_, RII_indirect_ and RII_sum_, respectively, were the response variables, the receiver species, the donor species, the selection history and all possible interactions were the explanatory variables, the species composition was the random term. Type-I analyses of variance (ANOVA) were performed to test the hypotheses. Afterwards, post-hoc analyses were conducted to test if there are significant differences between selection histories and/or culture × selection history (estimated marginal means, R package *emmeans* (Lenth, 2021)). For direct, indirect and total interaction intensities, estimated marginal means of the interaction receiver species × donor species × selection history were tested against zero (one-sample t-test). The interaction intensities significantly different from zero were used in the species interaction networks (**Fig. S2**).

Intransitivity and transitivity of interactions between three species were defined according to Gallien et al. (2018) (**Table 1**). An additional denotation was introduced to describe interactions that did not include all three species (i.e. pure pairwise). Transitivity/intransitivity was only described on species interaction network with RII_sum_ (**Fig. 4e-f**).

**Table 1.** Interaction denotations used in this study.

| denotation of interaction | interaction |
| --- | --- |
| pure pairwise | A>B, A>C, B>C |
| transitive (“hierarchical”) | A>B>C and B>C |
| weak intransitive | A>B>C=A |
| intransitive (“rock-paper-scissor”) | A>B>C>A |

To test for a relationship between net interaction intensity (estimated marginal means, **Fig. 3**) and the sum of all direct and indirect interaction intensities a species received (**Eq. 5**; estimated marginal means, **Fig. S4 and S5**), a simple linear regression was conducted (RII_net_ ~ RII_calc_).

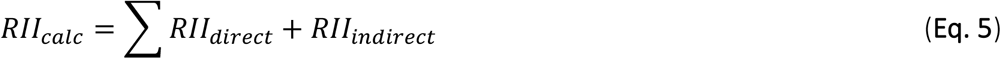

**Fig. 3.**
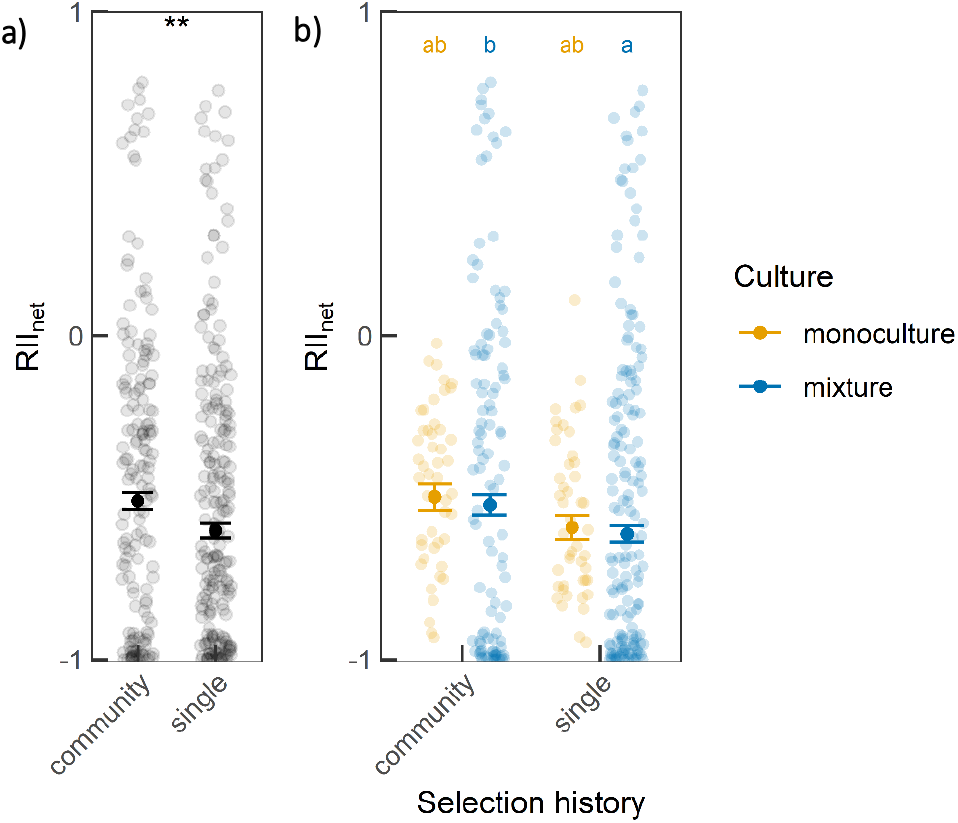
Net interaction intensity (RII_net_) of plants with either *community* or *single* selection history. (**a**) RII_net_ across all species. Shown are the single data points, the estimated marginal means ± standard error and the results from the contrast analysis between selection histories (asterisk). Significance levels correspond to *: P<0.05, **: P<0.01, ***: P<0.001. (**b**) RII_net_ across all species but between culture and selection history. Shown are the single data points, the estimated marginal means ± standard error and the grouping according to the contrast analysis between culture × selection history.

**Fig. 4.**
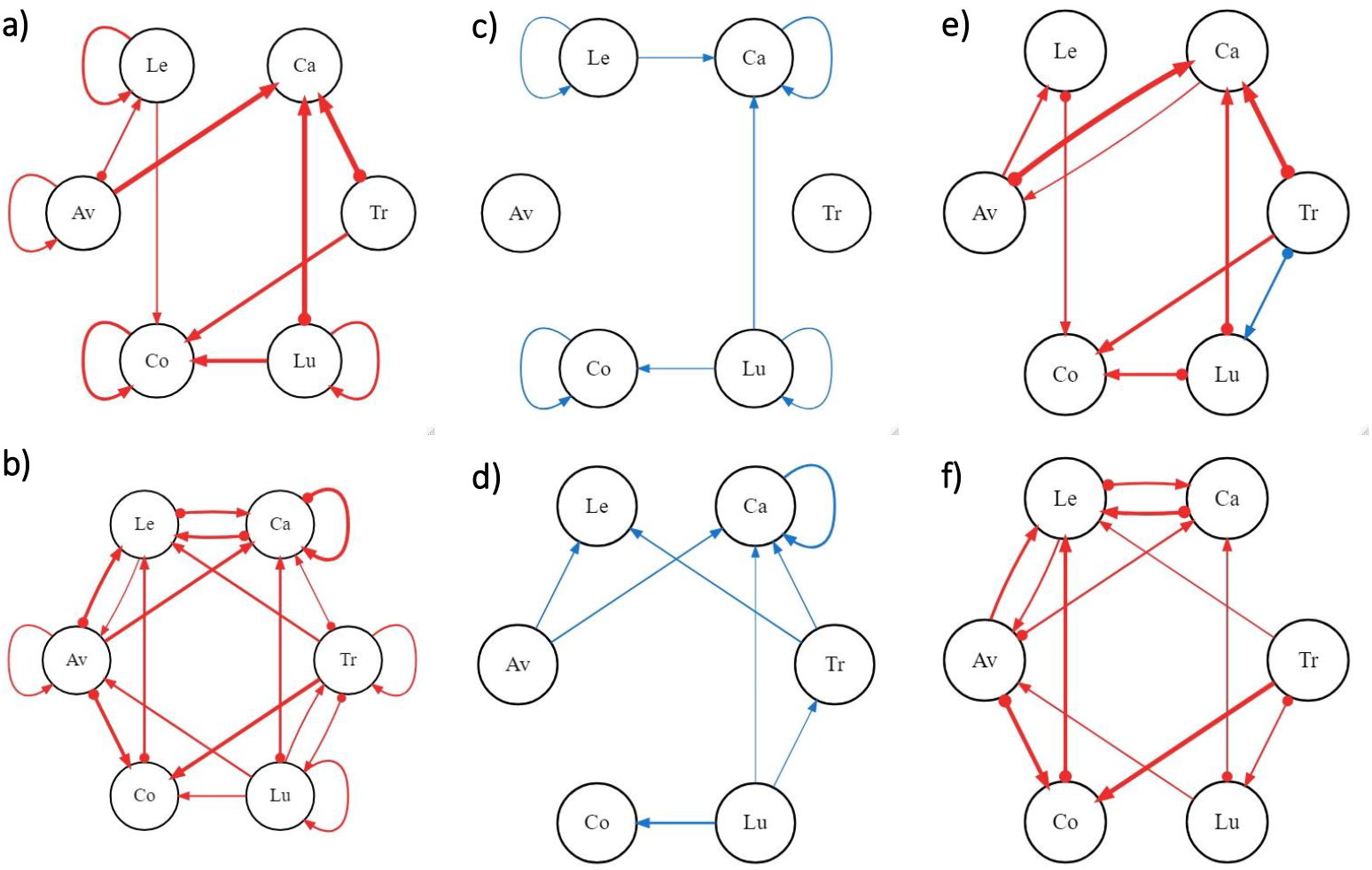
Direct (**a-b**), indirect (**c-d**) and total interactions (**e-f**) in communities that consisted of plants with either *community* (**a**, **c**, **e**) or *single* selection history (**b**, **d**, **f**). Blue arrows represent positive, red negative interactions. Shown are only the interaction intensities (estimated marginal means) that were significantly different from zero. Significant differences between the selection histories are indicated with a dot on the arrow tail (compare to S4, S5 and S6). Arrow thickness correspond to the interaction intensity (stronger positive or negative interaction intensities are thicker). Species abbreviations correspond to Av=oat, Tr=wheat, Le=lentil, Lu=lupin, Ca=camelina and Co=coriander.

## Results

### Interaction intensities in communities with and without coexistence history

RII_net_ was significantly less negative in *community* than *single* selection history across all species (**Fig. 3a**, **Table 2**). Nevertheless, this effect was only present in mixtures whereas in monocultures the same, but weaker (not significant) trend was visible (**Fig. 3b**). Especially oat, lentil and lupin in mixtures and coriander from monoculture plots experienced a less negative RII_net_ in *community* selection history (**Fig. S3**). For camelina from mixture plots, RII_net_ was significantly but only slightly less negative in *single* than *community* selection history. In general, for most plants, RII_net_ was negative and very strong. Only lupin grown in mixture experienced facilitative effects – these positive effects were even more pronounced in *community* selection history. The culture alone had a large impact especially on RII_net_ of lupin, camelina and coriander (**Fig. S3**). This indicates that intra- and interspecific interactions had varying effect on species RII_net_.

**Table 2.** Type-I analysis of variance of the response variables net (RII_net_), direct (RII_direct_), indirect (RII_indirect_) and total interaction intensity (RII_sum_). For RII_net_, the explanatory variables were species, culture, selection history and all possible interactions between the three variables. For the others (RII_direct_, RII_indirect_ and RII_sum_) the explanatory variables were receiver species, donor species, selection history and all possible interactions between the three variables. The random term was species composition. DF: degrees of freedom; DenDF: degrees of freedom of error term; F: probability distribution; P: error probability. P-values in bold are significant at α = 0.05.

|  | DF | DenDF | F | P |
| --- | --- | --- | --- | --- |
| $RII_{net}$ | | | | |
| species | 5 | 42.61 | 141.90 | **<0.001** |
| culture | 1 | 7.55 | 0.00 | 0.972 |
| selection history | 1 | 332.48 | 12.54 | **<0.001** |
| species × culture | 5 | 13.17 | 47.93 | **<0.001** |
| species × selection history | 5 | 328.66 | 3.92 | **0.002** |
| culture × selection history | 1 | 329.72 | 0.02 | 0.882 |
| species × culture × selection history | 5 | 326.54 | 4.57 | **<0.001** |
| $RII_{direct}$ | | | | |
| receiver species | 5 | 53.15 | 18.09 | **<0.001** |
| donor species | 5 | 67.67 | 7.34 | **<0.001** |
| selection history | 1 | 448.64 | 2.40 | 0.12 |
| receiver species × donor species | 19 | 59.17 | 1.52 | 0.111 |
| receiver species × selection history | 5 | 558.62 | 5.29 | **<0.001** |
| donor species × selection history | 5 | 558.19 | 1.58 | 0.164 |
| receiver species × donor species × selection history | 19 | 557.51 | 3.78 | **<0.001** |
| $RII_{indirect}$ | | | | |
| receiver species | 5 | 46.42 | 4.56 | **0.002** |
| donor species | 5 | 57.82 | 3.87 | **0.004** |
| selection history | 1 | 426.46 | 0.86 | 0.354 |
| receiver species × donor species | 19 | 46.61 | 2.46 | **0.006** |
| receiver species × selection history | 5 | 545.88 | 1.53 | 0.178 |
| donor species × selection history | 5 | 545.44 | 1.32 | 0.252 |
| receiver species × donor species × selection history | 19 | 544.58 | 1.76 | **0.025** |
| $RII_{sum}$ | | | | |
| receiver species | 5 | 55.39 | 9.98 | **<0.001** |
| donor species | 5 | 46.08 | 3.11 | **0.017** |
| selection history | 1 | 467.94 | 0.95 | 0.330 |
| receiver species × donor species | 19 | 30.42 | 3.06 | **0.003** |
| receiver species × selection history | 5 | 543.21 | 8.00 | **<0.001** |
| donor species × selection history | 5 | 543.10 | 1.64 | 0.147 |
| receiver species × donor species × selection history | 19 | 542.84 | 6.81 | **<0.001** |

Both RII_direct_ and RII_indirect_ were affected by selection history, but the effect depended on the receiver and donor species (**Table 2**). Subsequent post-hoc analyses revealed that in most cases, negative RII_direct_ were reduced in *community* selection history (**Fig. S4**). In the two cereals oat and wheat, the selection history had no effect on interaction intensity. The receiver species lentil and lupin always showed reduced negative RII_direct_ (i.e. competition) in *community* selection history. In camelina, this trend depended strongly on the donor species. For RII_indirect_, significant difference between selection history were only present in lentil and camelina – and also only imposed by the donor species oat and wheat (**Fig. S5**). In all these cases, *single* selection history showed increased positive RII_indirect_ (i.e. facilitation).

RII_sum_ (sum of direct and indirect interactions) showed a similar pattern like RII_direct_ (**Fig. S6**, **Table 2**). In some cases, the RII_indirect_ counteracted the effect of selection history on RII_direct_ (e.g. in receiver-donor pairs lentil-wheat and camelina-camelina). In other cases, RII_indirect_ enhances selection history effects on RII_direct_. These include the receiver-donor pairs camelina-wheat, coriander-lentil and coriander-lupin. In all these cases, RII_indirect_ especially reduced negative direct interactions in *single* selection history.

### Species interaction networks

The number of significant intraspecific direct interactions was largely affected by selection history (**Fig. 4a-b**). Direct interactions were generally fewer in *community* than in *single* selection history. In *community* selection history, coriander and camelina received most of the (negative) direct interactions – oat, wheat and lupin were the species that imposed most of the interactions (**Fig. 4a**). In comparison to *community* selection history, it was additionally lentil in the *single* selection history that received several negative direct interactions (**Fig. 4b**). Moreover, in *single* selection history, oat was also more frequently involved in receiving and imposing negative direct interactions. Reciprocal negative interactions that were present in *single* selection history do not appear in community selection history (e.g. in oat-lentil, lentil-camelina and wheat-lupin). Contrary to intraspecific direct interactions, the number of interspecific interactions was generally less affected by selection history (**Fig. 4a-b**).

Significant indirect interactions were generally fewer than direct interactions, always positive and had a similar extent in both selection histories (**Fig. 4c-d**). Surprisingly, positive indirect interactions especially in monocultures seemed very effective in mitigating negative direct interactions. In all monocultures and both selection histories, previously negative direct interactions were mitigated by indirect interactions (**Fig. 4e-f**). Other than that (especially in intraspecific interactions), indirect interactions could not counteract the strong negative direct interactions. This can mainly be explained by the presence of stronger negative direct interactions between species than within species.

Intransitivity and transitivity of interactions was greatly influenced by selection history (**Fig. 4e-f**, **Table S2**). In *community* selection history, from all the possible interaction between three species, most interactions were pure pairwise (e.g. oat>lentil, lupin>camelina; **Table S2**). There was only one weak intransitive interaction present between oat, lentil and coriander (i.e. oat>lentil>coriander=oat). In contrast, interactions in *single* selection history were mostly weak intransitive. Additionally, two transitive interactions were also present in *single* selection history (i.e. lupin>oat>camelina and lupin>camelina, wheat>coriander>lentil and wheat>coriander; **Table S2**).

## Discussion

With the method applied here to estimate direct and indirect interaction intensities between plants, we were able to study how co-adaptation affects these interactions in an agricultural system. We found that co-adaptation reduced overall competition and negative direct interactions between species especially in more diverse plant communities. Co-adaptation had a substantial impact on the interplay in the species interaction network and shifted mainly direct interactions among plants.

### The importance of direct and indirect interactions in community adaptation

In this study, we found evidence for reduced competition after co-adaptation of species in a community (**Fig. 3a**). This finding is in line with a study that also found reduce competition in communities with a common coexistence history (Stefan et al., 2022). Furthermore, community adaptation reduced competition particularly between species (i.e. interspecific interactions) whereas competition among species (i.e. intraspecific interctions) were not affected (**Fig. 3b**). Additionally, facilitative effects of the community on lupin were only present in mixtures and more pronounced upon community adaptation (**Fig. 3b**). This confirms that species diversity is crucial for the evolution of facilitation among plant species (Schöb et al., 2018). Besides, species diversity is also suggested to enhance indirect interactions and mitigate negative interactions (Aschehoug and Callaway, 2015). The net effects of the community on the species were also reflected in the direct and indirect interactions the species received. In fact, the sum of direct and indirect interaction intensities (as quantified with RII in our study) was a good predictor of the net effects (**Fig. S7**). This suggests that RII is a suitable measure to partition net effects into direct and indirect interactions.

In our system, direct interactions were strong and always negative, which implies strong competition among the six crop species. This can probably be explained by the high plant density within the small plots. Furthermore, direct interactions were calculated with the plant biomass in 3×3 Latin squares (9 individuals) and with the plant biomass in 2×2 Latin squares (4 individuals) as control. The additional species (which imposed the direct interaction) in the 3×3 Latin squares was always accompanied by a higher plant density. Consequently, significant direct interactions were exclusively negative, as expected (Miller, 1994; Vandermeer, 1990).

Beside that community adaptation significantly reduced competitive effects in the community (**Fig. 3**), it also decreased the intensity and number of negative direct interactions (**Fig. S4**, **Fig. 4**). As far as we know, this is the first study that could demonstrate a positive effect of community adaption on competitive direct interactions. Nevertheless, direct interactions were generally more dominant than indirect interactions and the facilitative effects of indirect interactions could not counteract the strong competitive effects of direct interactions (**Fig. 4**). Consequently, indirect interactions were rather insignificant for community adaptation in our study system. There is both theoretical (Lawlor, 1979; Levine et al., 2017) and empirical evidence (Cuesta et al., 2010; Michalet et al., 2015; Schöb et al., 2013) that underpin the importance of indirect interactions for plant coexistence and community structure. Additionally, indirect interactions are suggested to have evolutionary consequences on communities (Guimarães et al., 2017; Wootton, 1994). Yet, empirical evidence about the importance of indirect interaction for adaptation of plants in plant communities is lacking. This might be due to the fact the indirect interactions are generally hard to measure as they are not as apparent as direct interactions (Strauss, 1991).

### The effect of community adaptation on transitive and intransitive interactions

In a three species interaction network, transitive interactions occur when one species is more competitive than the other two and one is the least competitive (A>B>C and B>C, **Fig. 1d**, **Table 1**). On the other hand, intransitive interaction between three species arises when each species is more competitive than one of the others (A>B>C>A, **Fig. 1c**, **Table 1**). However, “strong” intransitive interactions are hardly found in nature (Soliveres and Allan, 2018). Thus, it was suggested that weak forms of intransitivity can also occur (“weak” intransitive interaction, A>B>C=A, **Table 1**) (Gallien et al., 2018). In our system, community adaptation reduced the number of (weak) intransitive interactions (**Table S2**). In most cases, these weak intransitive interactions were diminished to pure pairwise interactions where some species (mostly wheat, lupin and partly oat) were more competitive than other species (mostly lentil, camelina and coriander). Furthermore, transitive interactions were only present in communities without coexistence history (e.g. lupin>oat>camelina and lupin>camelina). Upon community adaptation hierarchical negative interactions were diminished to pure pairwise interactions.

Intransitive interactions are also affected by the heterogeneity of the environment (Allesina and Levine, 2011). In less fertile and drier environments, intransitive interactions are more common (Soliveres et al., 2018, 2015). This has large implications for cropping systems, which are usually highly productive (high fertiliser inputs) and very homogeneous. Under these conditions, intransitive interactions are less likely to exist and evolve (Gallien et al., 2018; Soliveres et al., 2018).

Intransitive competition can certainly be important among interacting crop species (i.e. in intercropping) as it reduces competitive differences among species (Gallien et al., 2017). However, an intransitive interaction that is very competitive probably also does not promote community productivity, which is the main goal in agricultural systems. Thus, the breeding of cooperative plants is essential to enhance agricultural productivity (Weiner, 2019; Weiner et al., 2017; Wuest et al., 2022). In our system, community adaptation increased competitive ability especially in lupin grown in mixture – and competitive ability of others such as camelina and coriander in mixtures decreased. Consequently, community adaptation did not promote cooperativeness of the crops, but rather resulted in an imbalance of competitive ability among species (i.e. asymmetric competition (Weiner, 1990)). Nevertheless, this does not imply that community productivity is lower upon co-adaptation – but since this was examined in this study, no conclusions can be drawn about this. Though, it would be interesting to link interactions among plants to community productivity.

### Conclusion

In this study, we investigated how community adaptation of crop species affects net, direct and indirect interaction intensities and how transitivity and intransitivity of interactions were shifted upon community adaptation. We demonstrated the competitive net and direct interaction intensities were mitigated upon community adaptation. Even though indirect interactions were generally facilitative, they could not counteract the strong competitive effects of (intraspecific) direct interactions in our system. Facilitation was only found in lupin grown in mixtures and was also more pronounced upon community adaptation. Moreover, we observed that community adaptation diminished (weak) intransitive interactions to pure pairwise competitive interactions.

## Acknowledgements

We thank Sandra González Sánchez, Francicso Ordiales Rosado, Jesús López-Angulo, Pilar Hurtado and Markus Bittlingmaier for their support in the field and Elisa Pizarro Carbonell n from the Aprisco association for using their facilities. This work was funded by the ETH Zurich Research Grant ETH-28 19-2. The authors declare that they have no conflicts of interest.

## Authors contribution

AS planned and conducted the experiment, analysed the data, and wrote the manuscript. CS obtained funding, planned the experiment and gave advise during the experiment, analysis and manuscript writing.

## Data availability statement

The data used in this study is available on Zenodo: https://doi.org/10.5281/zenodo.7621220.

