## Supplementary 1 for "Transgenerational coexistence history attenuates negative direct interactions and strengthens facilitation"

**Table S1**. Overview of the different planting schemes. The shading represents the different selection histories, the shapes (circle, triangle and square) are different species.

| **Planting scheme** | **Culture** | **Selection history** | |
| --- | --- | --- | --- |
|  |  | **community** | **single** |
| Single plant  (1 individual) |  | 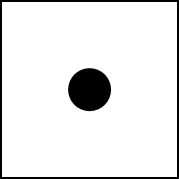 | 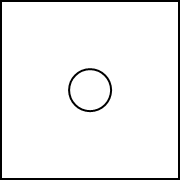 |
| 2x2 Latin square  (4 individuals) | monoculture | 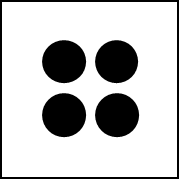 | 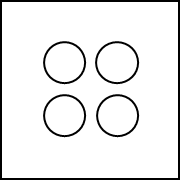 |
|  | mixture | 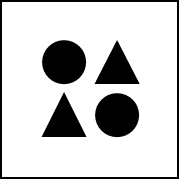 | 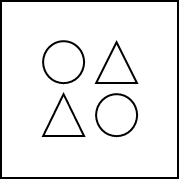 |
| 3x3 Latin square  (9 individuals) | monoculture | 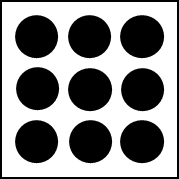 | 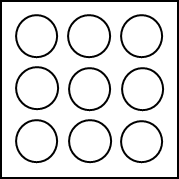 |
|  | mixture | 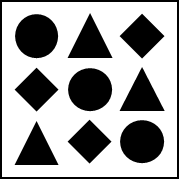 | 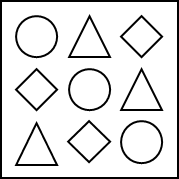 |

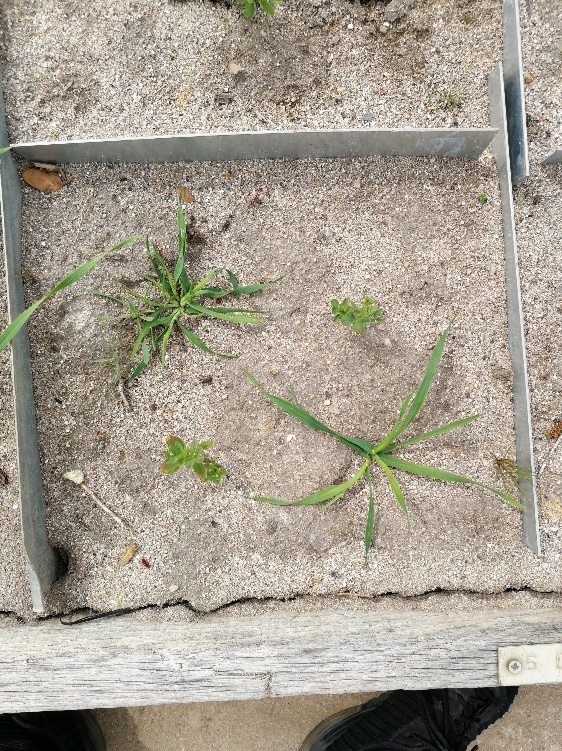

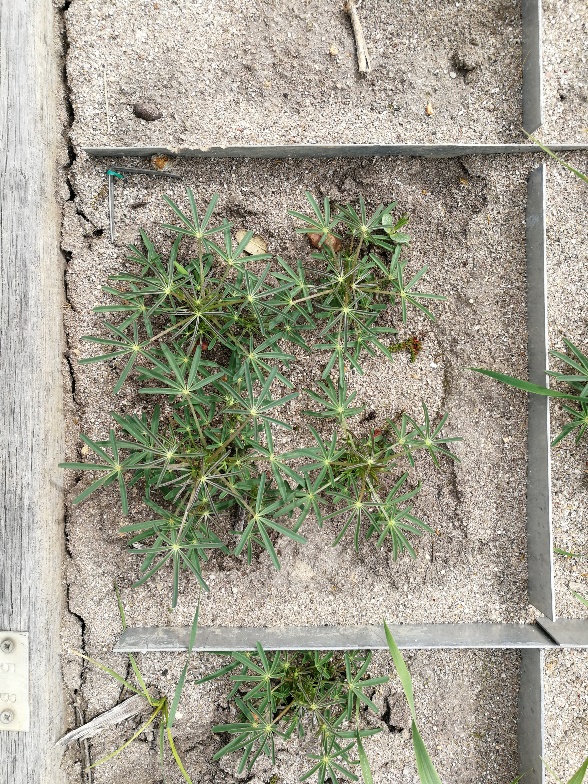

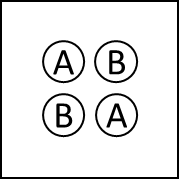

**a)**

**c)**

**b)**

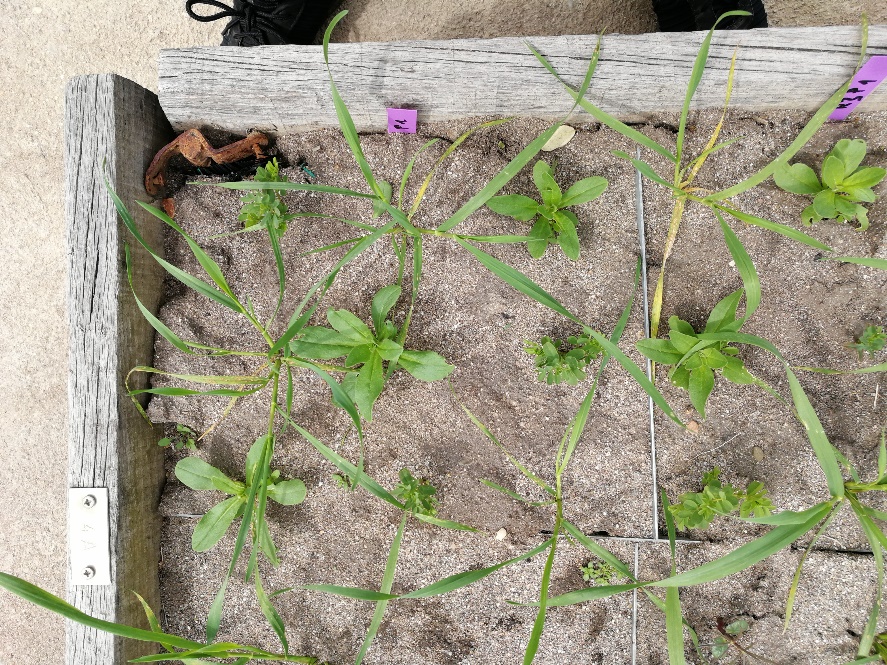

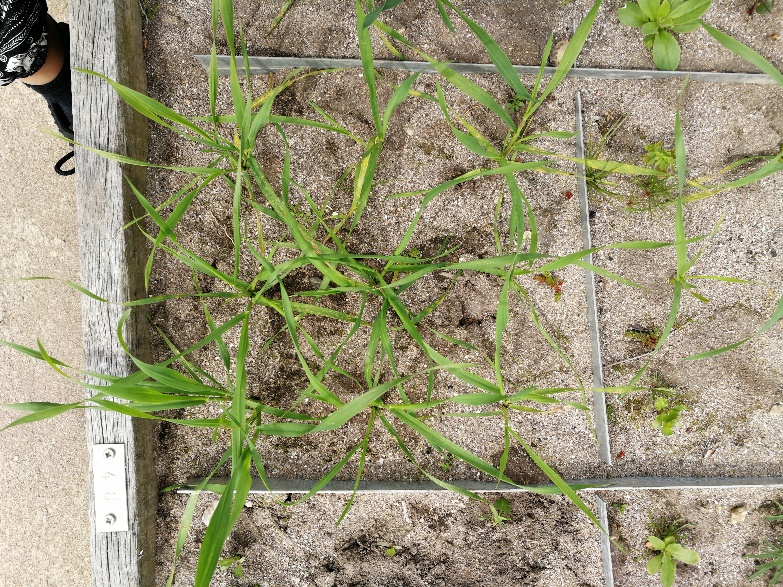

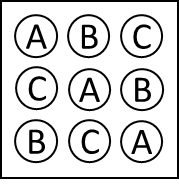

**f)**

**e)**
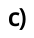
**)**

**d)**

**Fig. S1**. Examples of the planting scheme 2x2 Latin squares (**a-c**) and 3x3 Latin squares (**d-f**). (**a**) Mixture of wheat (position a) and lentil (position b). (**b**) Monoculture of lupin (position a and b). (**c**) The positions within the 2x2 Latin square. The individuals of the same species and selection history were always in the diagonal. (**d**) Mixture of camelina (position a), oat (position b) and lentil (position c). (**e**) Monoculture of wheat (position a, b and c). (**f**) The positions within the 3x3 Latin square.

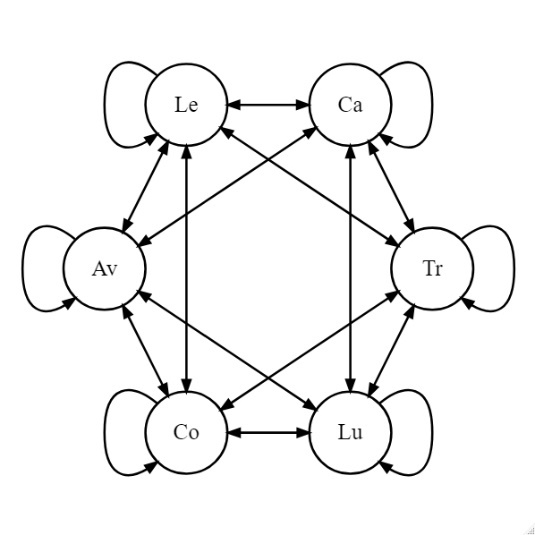

**Fig. S2**. All possible (direct and/or indirect) interactions in this study.

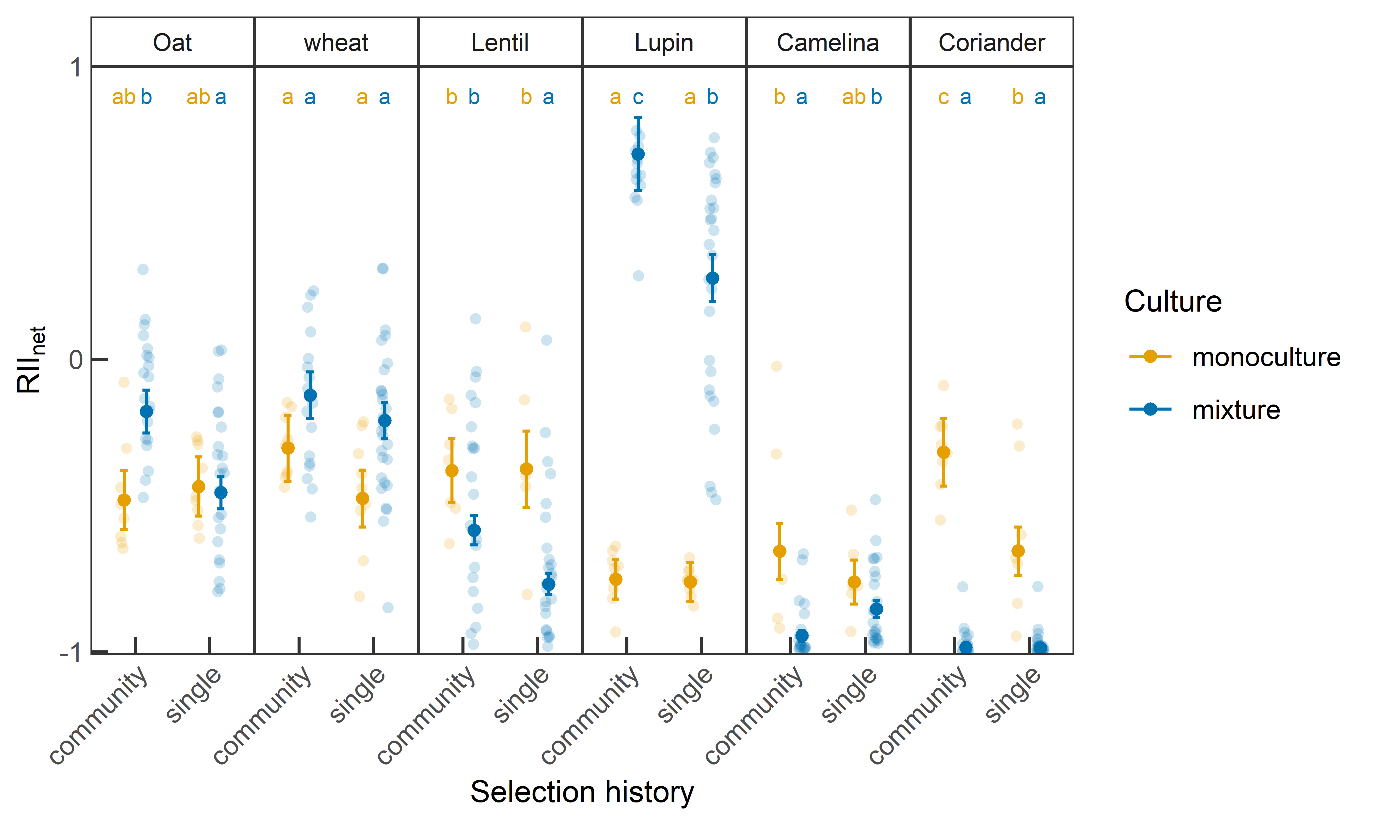

**Fig. S3**. Net interaction intensity (RII_net_) of the six species from either community or single selection history and grown in either monoculture (yellow) or mixture (blue). Shown are the single data points, the estimated marginal means ± standard error and the grouping according to the contrast analysis between culture × selection history within each species separately.

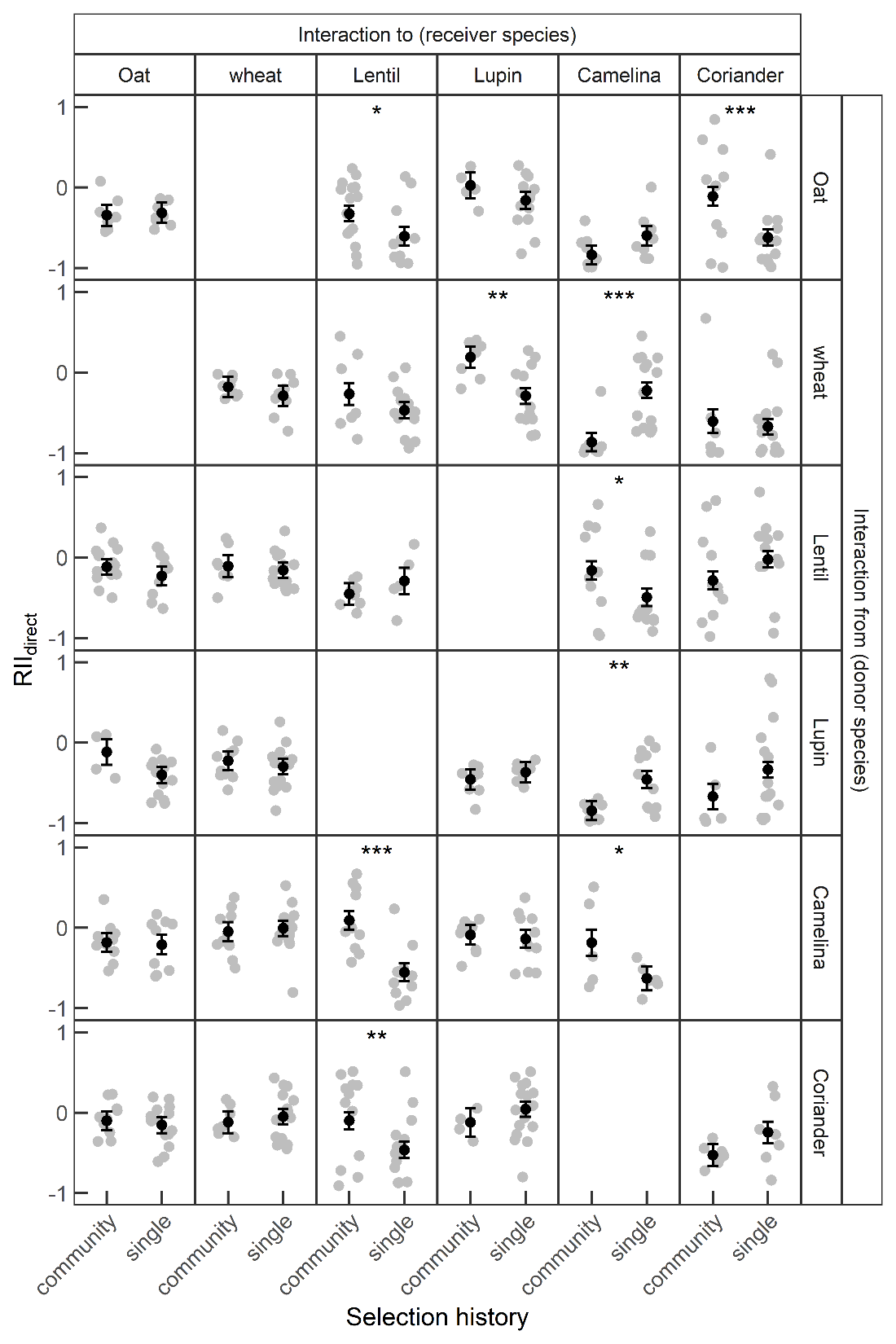

**Fig. S4**. Direct interaction intensity (RII_direct_) between receiver (interaction to) and donor species (interaction from)

of plants with either *community* or *single* selection history. Shown are the single data points, the estimated marginal means ± standard error and the results from the contrast analysis (asterisk). Significance levels correspond to *: P<0.05, **: P<0.01, ***: P<0.001.

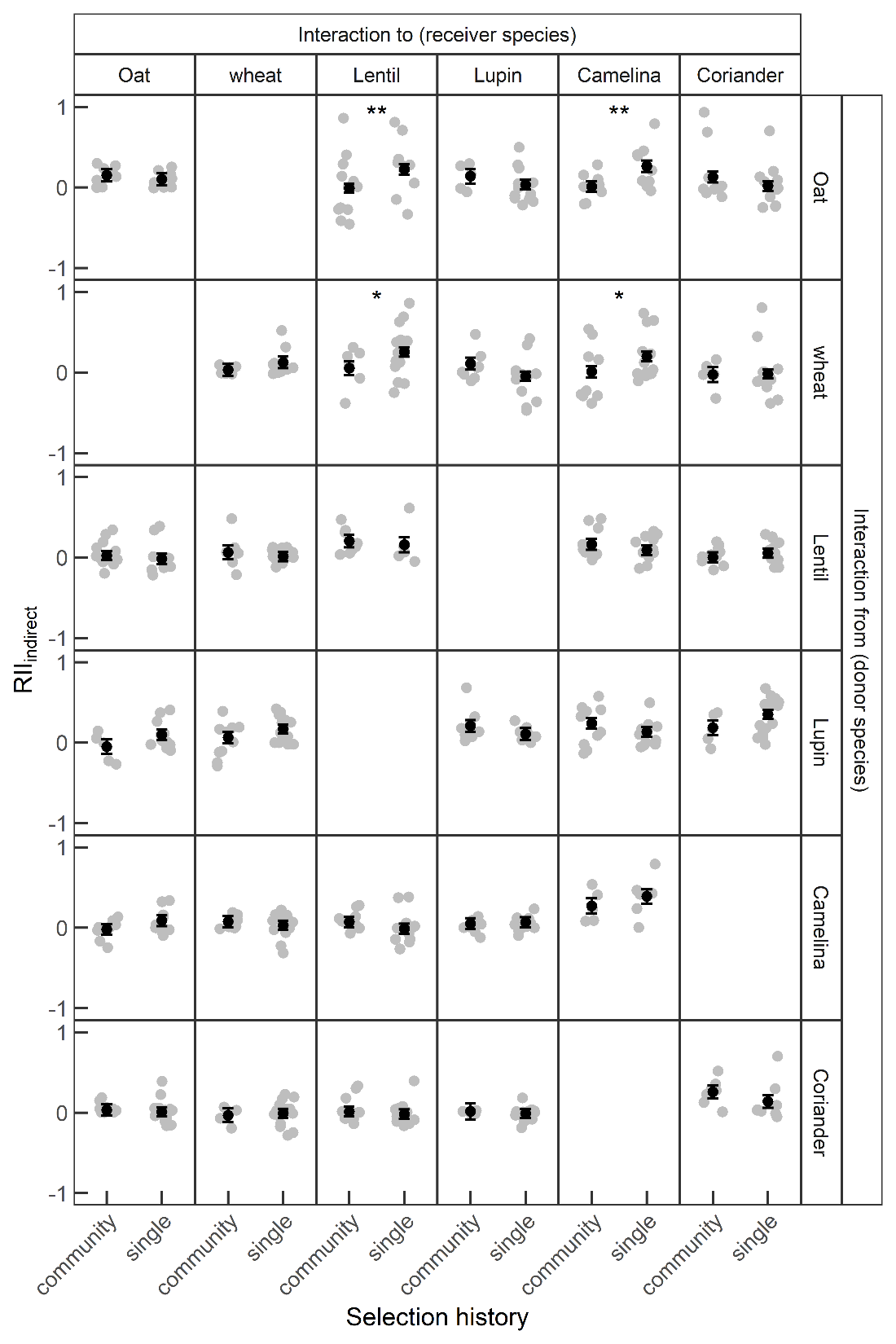

**Fig. S5**. Indirect interaction intensity (RII_indirect_) between receiver (interaction to) and donor species (interaction from) of plants with either *community* or *single* selection history. Shown are the single data points, the estimated marginal means ± standard error and the significant results from the contrast analysis (asterisk). Significance levels correspond to *: P<0.05, **: P<0.01, ***: P<0.001.

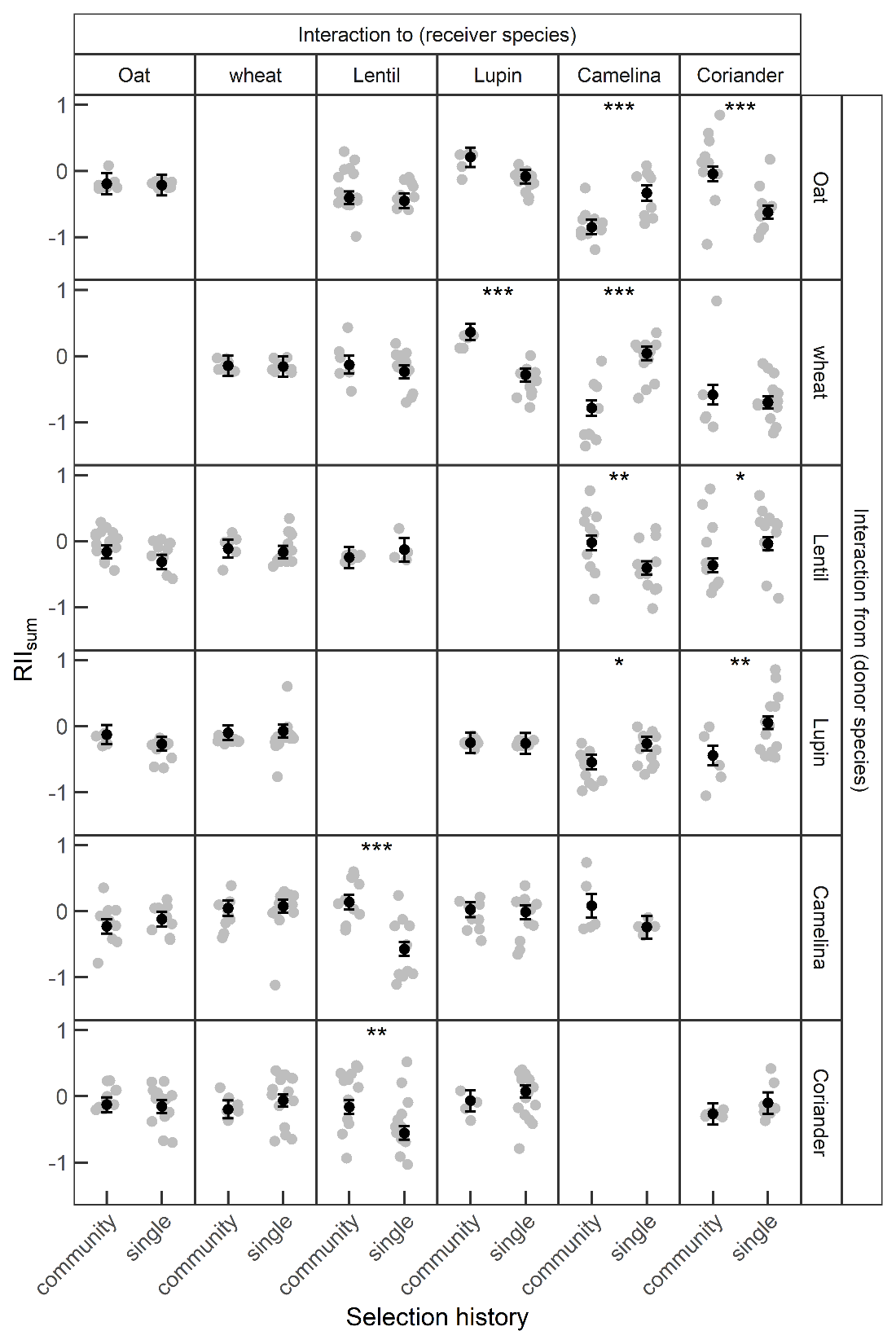

**Fig. S6**. Total interaction intensity (RII_sum_, sum of direct and indirect interaction intensities) between receiver (interaction to) and donor species (interaction from) of plants with either *community* or *single* selection history. Shown are the single data points, the estimated marginal means ± standard error and the significant results from the contrast analysis (asterisk). Significance levels correspond to *: P<0.05, **: P<0.01, ***: P<0.001.

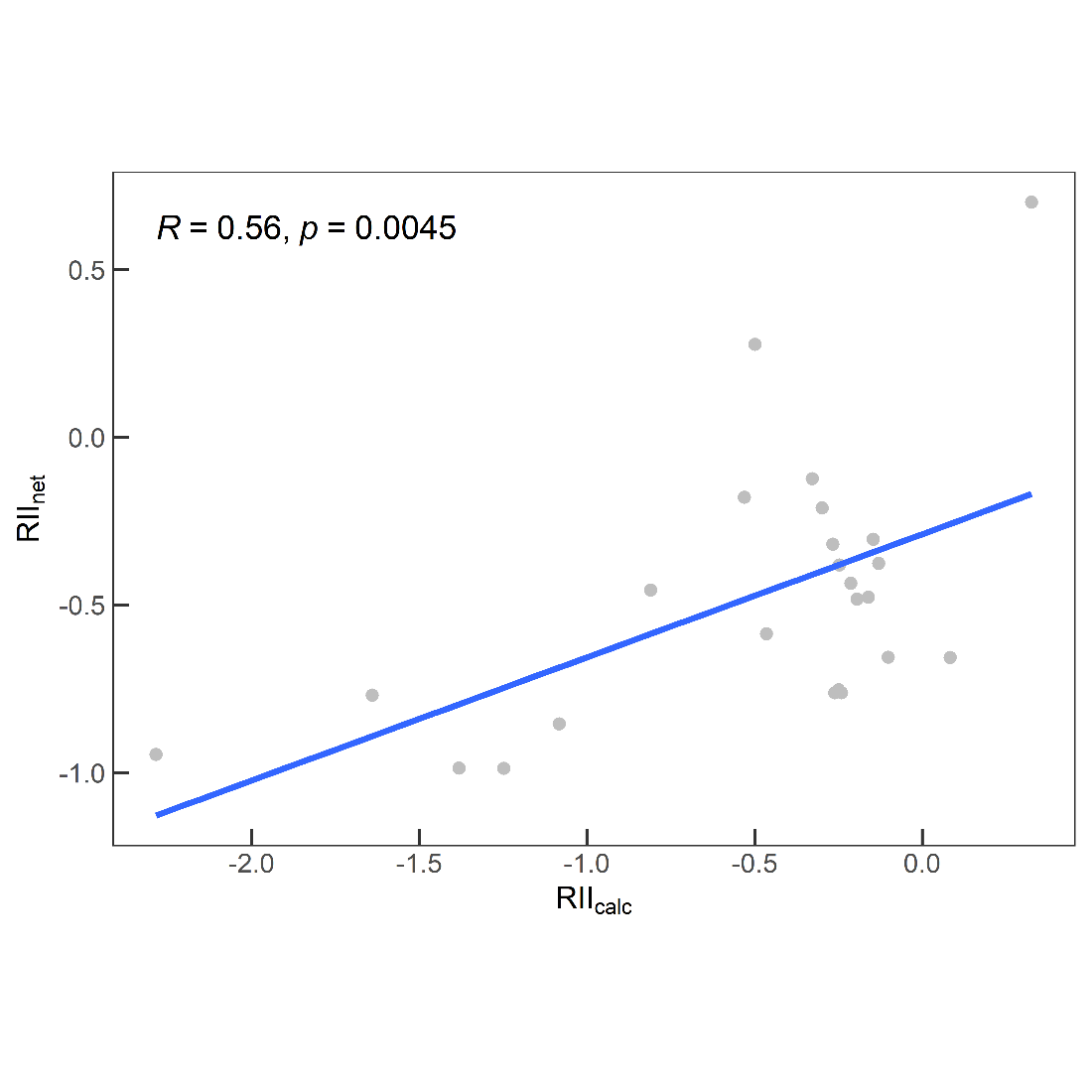

**Fig. S7**. Relationship between net interaction intensity (RII_net_, estimated marginal means, Fig. 3) and the sum of direct and indirect interaction intensities (RII_calc_, estimated marginal means, Eq. 5). Results from the Pearson correlation are indicated, and the regression line is drawn (blue line).

**Table S2**. Interaction denotations across all species compositions and for *community* and *single* selection history for total interaction intensity in Fig. 4e-f (RII_sum_, sum of direct and indirect interaction intensities). Species abbreviations correspond to Av=oat, Tr=wheat, Le=lentil, Lu=lupin, Ca=camelina and Co=coriander.

| selection history | species composition | interactions |
| --- | --- | --- |
| community | AvLeCa | pure pairwise (Av>Le, Av>=Ca) |
|  | AvLeCo | weak intransitive (Av>Le>Co=Av) |
|  | AvLuCa | pure pairwise (Av>=Ca, Lu>Ca) |
|  | AvLuCo | pure pairwise (Lu>Co) |
|  | TrLeCa | pure pairwise (Tr>Ca) |
|  | TrLeCo | pure pairwise (Tr>Co, Le>Co) |
|  | TrLuCa | pure pairwise (Lu>Ca, Tr>Ca) and facilitation (Tr🡪Lu) |
|  | TrLuCo | pure pairwise (Lu>Co, Tr>Co) and facilitation (Tr🡪Lu) |
| single | AvLeCa | weak intransitive? (Av>Le>=Ca>=Av) |
|  | AvLeCo | weak intransitive (Av>Co>Le>=Av) |
|  | AvLuCa | transitive (Lu>Av>Ca, Lu>Ca) |
|  | AvLuCo | weak intransitive (Lu>Av>Co=Lu) |
|  | TrLeCa | pure pairwise (Tr>Le, Le>=Ca) |
|  | TrLeCo | transitive (Tr>Co>Le, Tr>Le) |
|  | TrLuCa | weak intransitive (Tr>Lu>Ca=Tr) |
|  | TrLuCo | pure pairwise (Tr>Lu, Tr>Co) |
